# Active Function Learning

**DOI:** 10.1101/262394

**Authors:** Angela Jones, Eric Schulz, Björn Meder, Azzurra Ruggeri

## Abstract

How do people actively explore to learn about functional relationships, that is, how continuous inputs map onto continuous outputs? We introduce a novel paradigm to investigate information search in continuous, multi-feature function learning scenarios. Participants either actively selected or passively observed information to learn about an underlying linear function. We develop and compare different variants of rule-based (linear regression) and non-parametric (Gaussian process regression) active learning approaches to model participants’ active learning behavior. Our results show that participants’ performance is best described by a rule-based model that attempts to efficiently learn linear functions with a focus on high and uncertain outcomes. These results advance our understanding of how people actively search for information to learn about functional relations in the environment.

## Introduction

In everyday life, we constantly encounter situations where we have to make sense of rule-like, functional relations. How far can I drive if I put three gallons of gasoline in the tank? How much force should I apply to a football in order to hit the winning goal? How are different features of a food item related to its taste or calorie content? Historically, human function learning has often been treated as a passive information processing routine in which participants are sequentially confronted with different continuous stimuli (e.g., the length of a line) that are directly followed by a continuous response (e.g., the length of another line). In these studies, the input is either randomly determined or preselected by the experimenter, so that learning occurs incidentally rather than in a controlled, self-directed manner. This stands in contrast to many real-world scenarios, in which we *actively* seek out information, manipulating input features to observe and learn how they impact the outcome.

We present a multiple-feature function learning task to investigate how people actively select inputs to learn about their underlying functional relation with a continuous output. We use computational modeling techniques to gain insights into participants’ active learning. Considering both rule-based (linear regression) as well as non-parametric (Gaussian processes regression) approaches to active function learning, combined with different stepwise sampling strategies, we assess which active learning model best explains participants’ information selection. We find that behavior is best accounted for by a linear regression active function learning model that tries to learn about both uncertain and high outputs over time.

## Function Learning

Theories of function learning broadly coalesce around two different categories. *Rule-based theories* (Carroll, 1963) assume that participants learn explicit parametric representations, for example linear or polynomial functions. Even though rule-based theories can successfully predict function learning performance in some cases, they cannot account for all phenomena observed in human function learning. For example, they cannot explain why some rules, like linear functions, are easier to learn than others, like exponential functions (McDaniel & Busemeyer, 2005). Moreover, they fail to fully predict extrapolation performance (DeLosh, Busemeyer, & McDaniel, 1997) and are unable to learn a partitioning of the input space based on additional knowledge (Kalish, Lewandowsky, & Kruschke, 2004). On the other hand, *similarity-based theories* (e.g., Busemeyer, Byun, DeLosh, & McDaniel, 1997; DeLosh et al., 1997) assume that people perform function learning by associating inputs and outputs: if inputs **x** are paired with output *y*, then inputs similar to **x** should produce similar outputs to *y*. Busemeyer et al. (1997) formalized this intuition using a connectionist model (the *Associative-Learning Model;* ALM) in which inputs activate an array of hidden units representing a range of possible input values. Learned associations between the hidden units and the response units map the similarity-based activation patterns to output predictions. ALM broadly captures interpolation performance, but fails to explain some aspects of extrapolation and knowledge partitioning phenomena.

In order to overcome some of these problems, hybrid versions of the two approaches have been proposed (McDaniel & Busemeyer, 2005). Hybrid models of function learning contain an associative learning process that acts on explicitly-represented rules. One such hybrid is EXAM (*Extrapolation-Association Model;* DeLosh et al., 1997), which assumes similarity-based interpolation, but extrapolates using simple linear rules. EXAM captures the human tendency towards linearity, and predicts human extrapolations for a variety of functions. However, it does not account for non-linear extrapolation (Bott & Heit, 2004). Recently, Lucas, Griffiths, Williams, and Kalish (2015) put forward a theory of function learning based on *Gaussian Process* (GP) regression. GP regression is a non-parametric method to perform Bayesian regression and has been found to describe human functional judgments across a range of traditional function learning paradigms (Lucas et al., 2015). Moreover, GP models exhibit an inherent duality which makes them both a rule-based and a similarity-based model of function learning (see Lucas et al., 2015). As GP models have also been extended to account for exploratory behavior in function optimization tasks (Wu, Schulz, Speekenbrink, Nelson, & Meder, 2017), we will utilize them as candidate active learning models here.

## Active Learning

Active learning can be defined as a goal-directed behavior in which an agent tries to select information based on a measure of usefulness (Settles, 2009). Common metrics are the expected reduction of uncertainty, the extent of predictions’ improvement, or the maximization of future rewards (Nelson, 2005). Human active learning has been investigated in both adults and children (Nelson, Divjak, Gudmundsdottir, Martignon, & Meder, 2014; Ruggeri & Lombrozo, 2015; Ruggeri, Lombrozo, Griffiths, & Xu, 2016), and in several domains, including causal learning (Bramley, Dayan, Griffiths, & Lagnado, 2017; Lagnado & Sloman, 2004), categorization (Meder & Nelson, 2012), and control tasks (Osman & Speekenbrink, 2012). Active learning can often lead to better performance than passive learning (Markant, Ruggeri, Gureckis, & Xu, 2016). For instance, Lagnado and Sloman (2004) showed that participants actively intervening on a causal system performed better inferences about the underlying causal structure than yoked subjects who received identical information in a passive fashion. In category learning, Markant and Gureckis (2014) found that active learners sampled more along the line of the category boundaries and performed better than passive learners. Parpart, Schulz, Speekenbrink, and Love (2017) found that participants’ queries in a feature-based active learning task with binary outcomes were more in line with a weight-based strategy than a rank-based strategy.

## Experiment: Monster Top Trumps

Although there are several studies on active learning in different conceptual domains and experimental tasks, little is known about how humans search for information to learn about continuous functional relations. In the following, we first describe an active function learning experiment that we conducted and the obtained behavioral results. We then introduce different computational models for evaluating active learning strategies.

**Participants and Design** Participants were 98 adults, recruited from Amazon MTurk and tested online. They were randomly assigned to one of two learning conditions. In the *active condition* (*n* = 45) learners could freely choose which information they wanted to acquire, whereas in the *passive condition* (*n* = 53) they were presented with a random selection of instances. Average task duration was 14.61min (*SD* = 20.11). Mean reimbursement was $1.49 (*SD* = 0.29).

**Materials and Procedure** Participants played a browser-based card game, in which each card showed a different monster and values for its three features (Figure 1). The instructed goal was to learn the relationship between the monsters’ features (“friendly,” “cheeky,” and “funny”) and the number of “magic fruits” picked by each monster (criterion). Participants were told they would be rewarded for good performance in a subsequent test phase.

**Figure 1:**
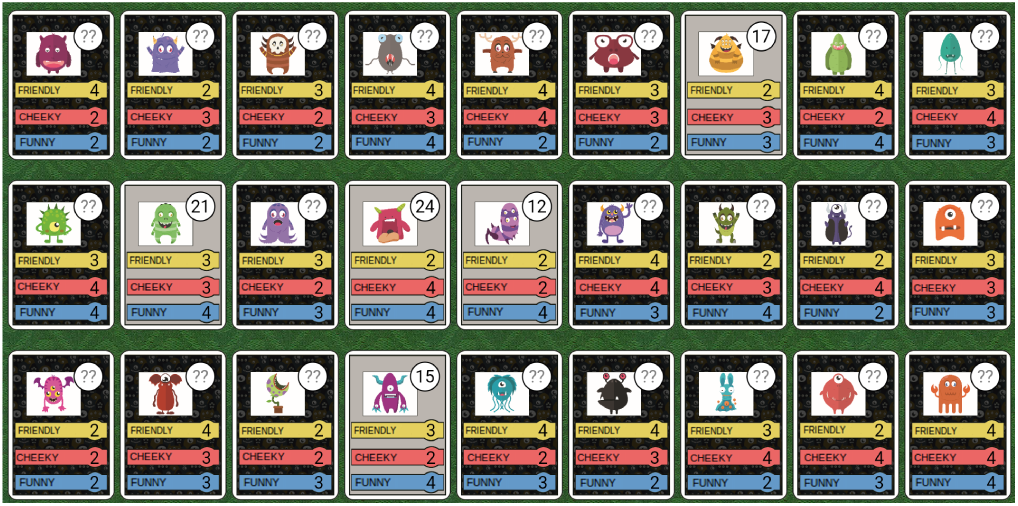
Screenshot of the multiple-feature function learning paradigm. The task was to learn the function relating the monsters’ features (“friendly,” “cheeky,” and “funny”) to the criterion (number of fruits picked, shown in top right corner of selected cards). The monster images and their assignment to each card were randomized.

The underlying function connecting the features to the criterion was the following:

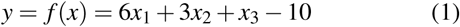

The weights for each dimension were decaying, to ensure that participants attended to all features in order to achieve good performance and could not easily use more simplistic strategies, such as tallying. For every participant, each of the features was randomly assigned to *x*_1_, *x*_2_ or *x*_3_. Feature values varied between one and five (inclusive), in increments of one. Participants were informed of this range before the learning phase.

**Learning Phase** Participants had to learn the function by step-wisely observing how a given monster’s feature values related to its criterion value. The learning phase card set consisted of 27 cards generated by factorially combining all feature values between 2 and 4, such that participants observed only a restricted part of the function. All 27 cards were displayed with the feature values visible but the criterion value hidden (Figure 1). In the active learning condition, participants freely selected cards to observe their criterion value. Participants in the passive learning condition selected randomly highlighted cards to see their criterion value. Thus, each participant received the same amount of information, but in one condition learners actively decided which data they wanted to observe, whereas the passive learners received randomly selected data points. All participants viewed the criterion value of 22 of the 27 cards. Once revealed, this value remained visible throughout the learning phase. However, they were not told exactly how many cards could be selected, such that the active learning group was incentivized to make the most informative selections from the beginning, while the passive group was encouraged to pay close attention to avoid missing any important information.

**Test Phase** The test phase consisted of two tasks: a *pair comparison* task and an *criterion estimation* task (order counterbalanced across participants). No feedback was given during the test phases.

In the pair comparison task, participants were shown eight card pairs whose feature values ranged between 1 and 5, such that these profiles contained both known and unknown feature values. For each pair they had to decide which monster had gathered more fruits; $0.04 was awarded for every correct selection. This task assessed how well they could judge the relative weights of each feature in the function they had to learn. For three of these trials, the card pairs were assembled such that one of the three features varied between cards while the values for the other two features were held constant. For the other five trials, card pairs were assembled so that the value of the first, second or last feature outweighed the combined value of the two other features on each card, so that this feature was the main determinant of the number of fruits collected.

The criterion estimation task consisted of three types of estimates: five recall trials, five interpolation trials, and eight extrapolation trials (18 cards in total; order randomized block-wise across participants). *Recall* cards consisted of five new monster cards with feature profiles for which participants had observed the criterion in the learning phase. *Interpolation* cards consisted of five new monsters with feature profiles corresponding to the five cards that had not been observed during the learning phase. Comparing performance on recall and interpolation trials enabled us to determine the relative contribution of memory versus inductively formed beliefs about the part of the function learners had been trained on. Features on the *extrapolation* cards had values of 1 or 5, corresponding to input values and combinations thereof that participants had not been trained on. For each card, estimates were given by moving a slider horizontally between 0 and 40 (in increments of 1) until it reached the desired criterion value. Performance was incentivized: estimates within 5 of the true criterion value were rewarded with $0.06; estimates within 10 were rewarded with $0.04; estimates within 20 were rewarded with $0.02; estimates further than 20 away from the criterion were not rewarded.

## Behavioral Results

### Learning Outcomes

**Pair Comparison** Overall, participants performed well in this task, with a mean accuracy of 77.14% (*SD* = 20%). Performance did not differ significantly between the active (*Mdn* = 75.0%) and passive (*Mdn* = 88.0%) conditions (Mann-Whitney-U, *U* = 1215.5, *p* = .87).

**Criterion Estimation** Performance in the criterion estimation task was assessed in terms of the estimation error (mean absolute deviation between participants’ estimates and the correct criterion value). Overall, performance did not differ between conditions (*Mdn_active_* = 5.60, *Mdn_passive_* = 6.38; *U* = 11232.5, *p* = .49). Performance was also evaluated separately for each trial type, aggregated across training conditions. There was a significant difference in estimation error between trial types (Kruskal-Wallis: *H*(*2*) = 16.83 adjusted for ties, *p* < .01). Post-hoc pairwise comparisons using the Dunn-Bonferroni method (Bonferroni-corrected) revealed that error on the extrapolation trials was significantly higher than on the interpolation (*p* < .05) and recall (*p* < .001) trials; there was no difference between interpolation and recall trials (*p* = 1.00).

We also examined differences in accuracy for each trial type according to the active vs. passive learning condition (Figure 2). Contrary to expectations, learning condition only marginally affected accuracy for recall (*Mdn_active_* = 4.2; *Mdn_passive_* = 4.6;U = 1206.0, *p* = .096), interpolation (*Mdn_active_* = 4.8; *Mdn_passive_* = 6.0; *U* = 1250.0, *p* = .69) or extrapolation trials (*Mdn_active_* = 7.1; *Mdn_passive_* = 8.4; *U* = 1289.5, *p* = .41).

**Figure 2:**
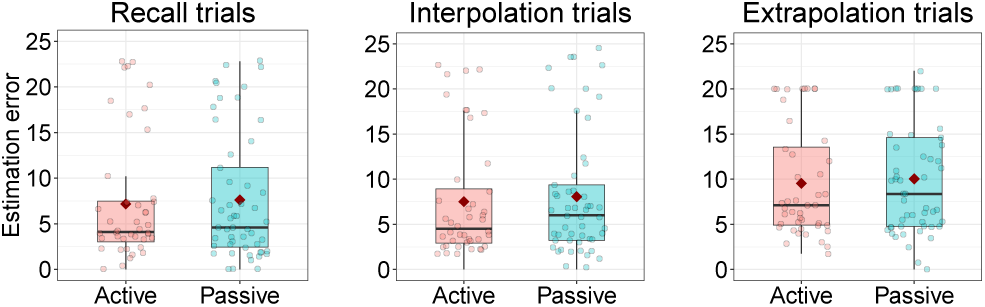
Estimation error (mean absolute difference) for the three criterion-estimation trial types, in each learning condition. Box plot whiskers indicate 1.5 IQR; diamonds show the mean, and lines show the median. Dots show individual data points.

### Active Learning Queries

We analyzed participants’ card selection in the active learning condition. Figure 3 shows how frequently participants selected different feature values for the top-most, the center, and the bottom-most feature. Participants tended to select extremes of the top-most feature during earlier trials, then switched focus and selected extreme values for the middle feature during intermediate trials, before then selecting extreme features for the bottom-most feature during later trials of the experiment. Comparing participants’ queries from the active condition against the (random) queries from the passive conditions, we found that queries within the first 10 trials significantly differed from random (Kolmogorov-Smirnov: *D* = 0.09, *p* < .001), whereas the last 10 queries did not (*D* = 0.036, *p* = .28). This suggests that the most systematic behavior occurred throughout the first few trials.

**Figure 3:**
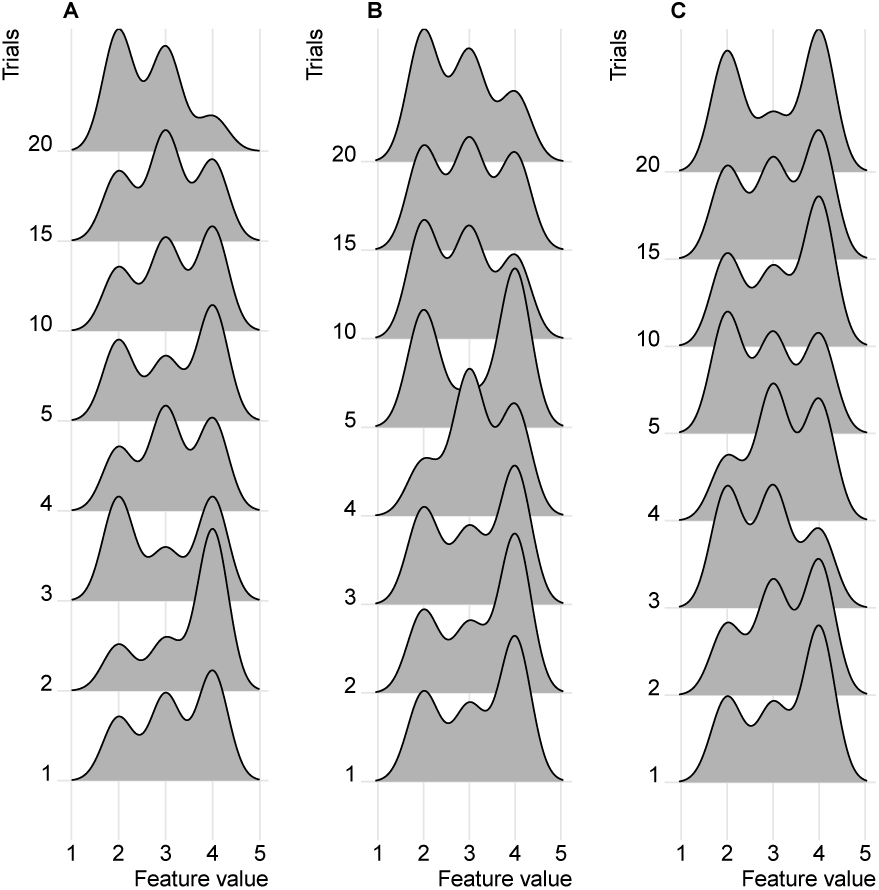
Proportions of chosen features across training trials. A indicates the top, B the middle, and C the bottom feature. Bandwidth of density estimates was set jointly to 0.27.

## Modeling active function learning

We compared two function learning models combined with different sampling strategies to see which model best accounted for participants’ active function learning behavior.

### Regression models for active function learning

We used two different regression models to describe participants’ active function learning behavior: a rule-based linear regression and a non-parametric GP regression.

**Linear Regression** A linear regression assumes that the outputs at time point *t* are a linear function of the inputs plus some added noise:

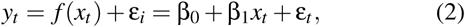

where the noise term *ε_t_* follows a normal distribution 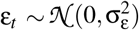 with mean 0 and variance 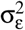. In matrix algebra, this can be written as

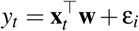

defining the vectors

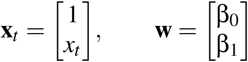

Within a Bayesian framework, we can compute the posterior distribution over the weights. Assuming a Gaussian prior 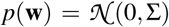 and a Gaussian likelihood 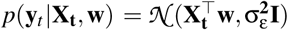, the posterior is

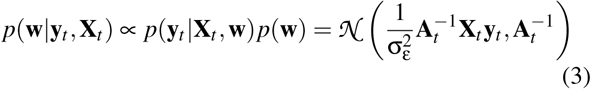

where 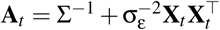. To predict a new output *y*_⋆_ at a new test point **x**_⋆_, one ignores the error term and only focuses on the expected value which is provided by the function *f*, predicting *f*_⋆_ = *y*_⋆_ − *e*_⋆_ = *f* (**x**_⋆_). In the predictive distribution of *f*_⋆_, the uncertainty regarding the weights is averaged out leading to:

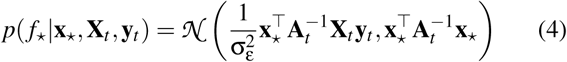

**Gaussian Process Regression** The other active function learning model is based on Gaussian Process (GP) regression. A GP regression is a non-parametric Bayesian way to model regression problems. It can theoretically learn any stationary function by the means of Bayesian inference (Schulz, Speekenbrink, & Krause, 2017). If 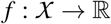 is a function over input space *χ* that maps to real-valued scalar outputs, then this function can be modeled as a random draw from a GP:

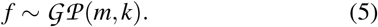

Here, *m* is a mean function that is commonly set to 0 to simplify computations. The kernel function *k* specifies the covariance between outputs.

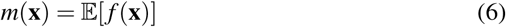

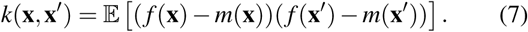

Conditional on observed data 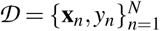, where 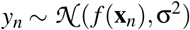 is a draw from the underlying function, the posterior predictive distribution for an input **x**\* is Gaussian with mean and variance:

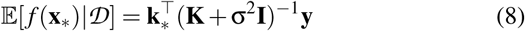

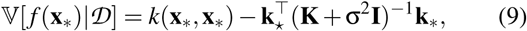

where 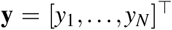, **K** is the *N* × *N* matrix of covariances evaluated at each pair of observed inputs, and **k**_*_ = [*k*(**x**_1_,**x**_*_),…,*k*(**x***_N_*, **x**_*_)] is the covariance between each observed input and the new input **x**_*_. The kernel function *k* encodes prior assumptions about the underlying function. Here, we used the *radial basis function* (RBF) kernel to model the underlying functional dependencies:

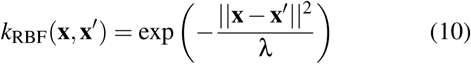

where λ governs the amount of correlation between **x** and **x**′.

### Active Sampling Strategies

Both function learning models generate predictions about the expected mean and associated uncertainties of outputs produced by different inputs. What is additionally required to guide active function learning is a sampling strategy that maps models’ predictions onto utilities. One simple sampling strategy is *uncertainty sampling*, which selects as the next point the one that is currently most uncertain, i.e., shows the highest predictive posterior standard deviation.

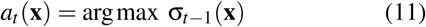

This drives down the uncertainty across the predictive space quickly. If the underlying parametric shape matches the used model of learning (for example, learning a linear function using linear regression), then this strategy results in theoretically fast learning. Moreover, this is the only active learning model that has proven guarantees when paired with Gaussian Process regression as it achieves at least a constant fraction of the maximum information gain possible (Krause, Singh, & Guestrin, 2008).

Another sampling strategy is *mean sampling* which selects as the next input point the one that currently promises to produce the highest output:

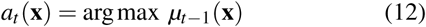

This strategy does not attempt to learn efficiently but rather learns about the function only serendipitously by the attempt to produce high outputs.

*Upper confidence bound sampling* tries to both reduce uncertainty and high outcomes by sampling the input that currently shows the highest upper confidence bound

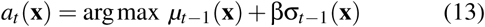
where β is a free parameter governing the extent to which participants sample uncertain options. UCB sampling will, on average, converge to both high knowledge about the underlying function and sampling the highest possible outcomes. It has recently been found to describe human behavior well in a function exploration-exploitation paradigm (Wu et al., 2017).

We also combined the linear regression model with two additional sampling strategies. The *weight-based sampling strategy* samples points that are expected to reduce the variance of the different regression weights maximally. The extent to which different observations reduce the weights’ variances is assessed by forward Monte Carlo sampling (i.e., sampling from the predictive distribution, concatenating that observation to the data, updating the Bayesian regression, assessing how much the weights’ variance has been reduced, and so forth). The *focused sampling strategy* is similar to the weight-based sampling strategy but instead of sampling points that promise to reduce the variance over all weights, only focuses on one weight at a time, namely the one that currently has the highest estimated variance. We combine the GP regression model with one additional sampling strategy, *active sampling as proposed by Cohn* (Cohn, Ghahramani, & Jordan, 1996). This sampling strategy selects as the next point the one that is expected to maximally reduce the predictive variance over the whole input space. This reduction is again assessed by using one-step ahead Monte Carlo samples as proposed by Gramacy and Lee (2009).

We did not assess any sampling strategy that considered more than one-step-ahead simulations. All models were assessed by submitting their predictions, generated by feeding participants’ observations up until trial *t* − 1 into a model and then predicting means and uncertainties for trial *t* over all trials, into a softmax function to convert the predicted utilities into choice probabilities

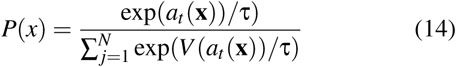
where *τ* s a free temperature parameter. For each participant we calculated a model’s 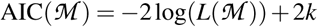 and standardized it using a pseudo-*R*^2^ measure as an indicator for goodness of fit, comparing each model 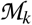 to a random model: 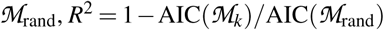. The free temperature parameter was optimized using differential evolution optimization (Mullen, Ardia, Gil, Windover, & Cline, 2009).

## Model Comparison Results

Figure 4 shows the model comparison results. Overall, the linear models described participants better than the GP models (averaged over mean, uncertainty and UCB; *t* (44) = 4.48, *p* < .001, *d* = 0.67). Only looking at the GP models, the uncertainty sampling strategy performed worse than chance*t*(44) = −9.90, *p* < .001, *d* = 1.47), indicating that participants did not sample the most uncertain options, and perhaps even avoided highly uncertain inputs. Furthermore, the UCB sampling strategy did not perform better than the mean sampling strategy when combined with a GP (*t*(44) = − 1.55, *p* < .13, *d* = 0.23). Only looking at the linear model, the UCB sampling strategy turned out to be best overall, directly followed by the mean sampling (linear model only; *t*(44) = 3.97, *p* < .001, *d* = 0.59) and the uncertainty reduction strategy (linear model only; *t*(*44*) *=* 5.24, *p* < .001, *d* = 0.78). The linear model paired with UCB-sampling performed marginally better than the GP paired with UCB sampling (*t*(44) = 1.78, *p* = .08, *d* = 0.27). Thus, the best overall model was a linear regression function learning algorithm paired with a UCB sampling strategy.

**Figure 4:**
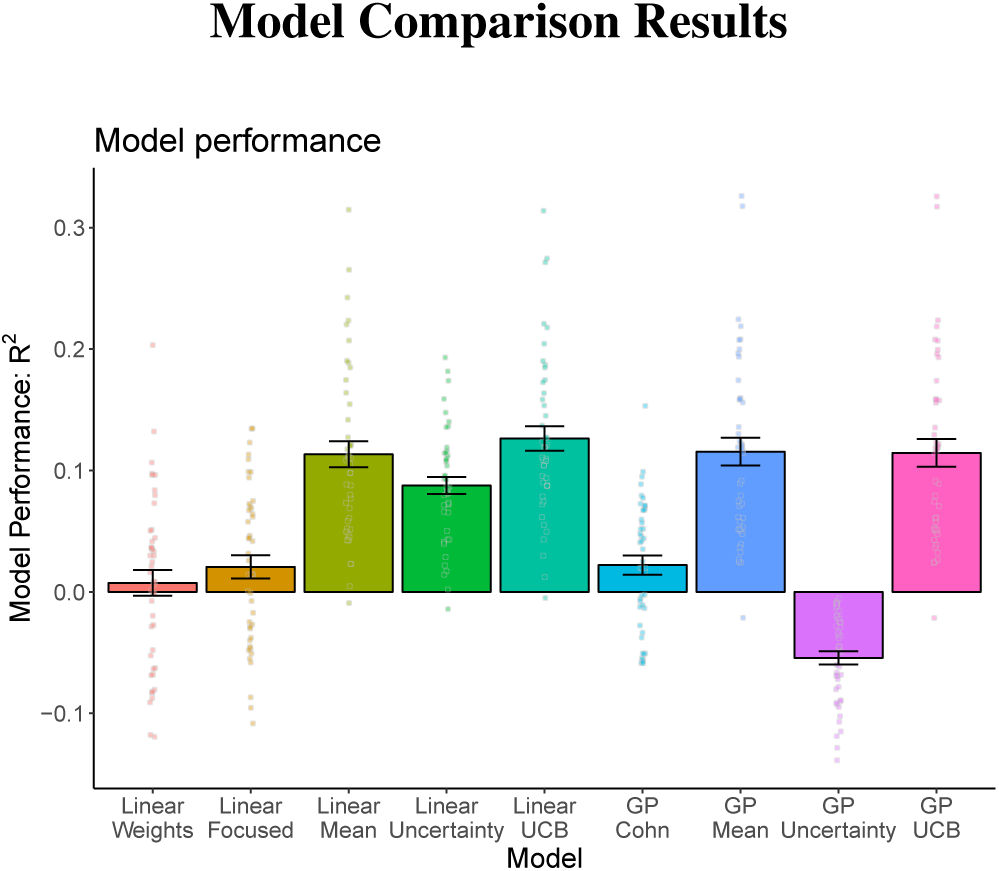
Results of model comparison. Error bars show the standard error of the mean. Dots show *R*^2^ for individual participants.

## Discussion and Conclusion

Function learning plays a crucial role across many domains, from completing basic visual patterns to predicting the future to gain rewards. We have developed and tested a novel paradigm to assess active function learning, exploring how participants step-wisely select inputs for which they want to observe outputs in order to learn about an underlying functional relation. To evaluate active learners’ behavior we compared linear and GP regression models combined with different sampling strategies. We found that the best overall model was a linear regression combined with an UCB sampling strategy. This means that participants adapted well to the overall task structure, which was set up to be linear, a finding which is in line with previous work showing that participants rely more on rule-based than similarity-based reasoning in simple linear environments (Hoffmann, von Helversen, & Rieskamp, 2016). Moreover, they sampled inputs to learn about both the overall shape of the function as well as the location of high outputs. Interestingly, this result mirrors recent evidence that participants solve the exploration-exploitation dilemma in reinforcement learning problems in a similar fashion (Wu et al., 2017), thereby hinting at the possibility of a universal sampling strategy underlying both information search and the search for rewards. Participants may not easily be able to turn off the exploitation part of their sampling strategy as they normally encounter a mix of exploration and exploitation problems in real life.

Surprisingly, our results showed no advantage of active information selection over passively observing outputs, possibly due to a ceiling effect after having observed 22 exemplars whose output remained on the screen during learning. In future studies, we intend to re-assess the possible advantage of active over passive function learning by further restricting participants’ sampling horizon. In fact, the considered models also predict no difference between conditions as they can learn the underlying function almost perfectly well within the 22 trials provided. Furthermore, it may be informative to add a condition where participants are informed of the search horizon, so that they can plan their search accordingly. In addition, as we have only investigated a linear function, follow-up studies should attempt to analyze the effect of nonlinear functions on participants’ active function learning behavior; this in turn could lead to an increased performance of the GP model, which can learn functions well across many parametric forms. Finally, we believe that experimentally mapping out developmental differences in active function learning between children and adults promises to open an informative window onto the interplay between generalization and self-directed sampling across the lifespan.

## Acknowledgments

ES is supported by the Harvard Data Science Initiative; BM is supported by grant ME 3717/2-2 as part of the DFG priority program “New Frameworks of Rationality” (SPP 1516). We thank Federico Meini for his invaluable programming skills.

